# Personal Mastery Attenuates the Association between Greater Perceived Discrimination and Lower Amygdala and Anterior Hippocampal Volume in a Diverse Sample of Older Adults

**DOI:** 10.1101/2024.01.12.575447

**Authors:** Michael A. Rosario, Razan Alotaibi, Alan O. Espinal-Martinez, Amara Ayoub, Aletha Baumann, Uraina Clark, Yvette Cozier, Karin Schon

## Abstract

There is limited research investigating whether perceived discrimination influences brain structures that subserve episodic memory, namely the hippocampus and amygdala. Our rationale for examining these regions build on their known sensitivity to stress and functional differences along the long-axis of the hippocampus, with the anterior hippocampus and amygdala implicated in emotional and stress regulation. We defined perceived discrimination as the unfair treatment of one group by a dominant social group without the agency to respond to the event. A potential moderator of perceived discrimination is personal mastery, which we operationally defined as personal agency. Our primary goals were to determine whether perceived discrimination correlated with amygdala and anterior hippocampal volume, and if personal mastery moderated these relationships. Using FreeSurfer 7.1.0, we processed T1-weighted images to extract bilateral amygdala and hippocampal volumes. Discrimination and personal mastery were assessed via self-report (using the Experiences of Discrimination and Sense of Control questionnaires, respectively). Using multiple regression, greater perceived discrimination correlated with lower bilateral amygdala and anterior hippocampal volume, controlling for current stress, sex, education, age, and intracranial volume. Exploratory subfield analyses showed these associations were localized to the anterior hippocampal CA1 and subiculum. As predicted, using a moderation analysis, personal mastery attenuated the relationship between perceived discrimination and amygdala and anterior hippocampal volume. Here, we extend our knowledge on perceived discrimination as a salient psychosocial stressor with a neurobiological impact on brain systems implicated in stress, memory, and emotional regulation, and provide evidence for personal mastery as a moderating factor of these relationships.

## Introduction

The hippocampus and amygdala are part of the medial temporal lobe (MTL) system and have been well-established for their exquisite sensitivity to the impact of chronic psychosocial and physical stress^1–3^. The hippocampus can be functionally and structurally delineated along its long-axis, with the anterior portion implicated in affective function and posterior portion pivotal to spatio-cognitive tasks^4, 5^. The hippocampus is directly structurally connected to the amygdala^6^, and work within a larger structural and functional network to regulate stress and emotions^2, 7, 8^. Recent work reported greater psychosocial stress correlating with alterations in hippocampal and amygdala volume in humans^3, 9^. Parallel findings in rodent and nonrodent models using foot shocks, chronic restraint, and social stress paradigms have emphasized the sensitivity of these two brain regions to chronic stress^1, 2, 8–12^. For example, chronic stress exposure results in dendritic pruning and synapse loss in the amygdala and hippocampus, as well as apoptosis of adult-born neurons in the dentate gyrus subregion of the hippocampus^1, 2, 8–12^. Of particular importance and currently understudied is the role that psychosocial stress, namely perceived discrimination, has on amygdala and hippocampal structure and its implications for and translation to human health and well-being.

Here, we define perceived discrimination as the experience of unfair treatment of one group by a dominant social group along both singular and intersectional identities, with limited or no agency to respond to the event^13–15^. A subjugated identity can be defined along many social axes (e.g., race, gender, sexual orientation, disability, age, class, etc.). Although these identities are social constructs, perceived discrimination has dire consequences for biopsychological health and well-being^14, 16–18^. More experiences of discrimination have been associated with increased depression and anxiety^19^ and higher inflammation^18, 20^. Within the last decade, perceived discrimination has been well-characterized through cross-sectional and longitudinal research to have negative consequences for neurocognitive health^21–26^. In healthy Black older adults, greater perceived discrimination was associated with poorer global cognition, episodic memory and perceptual speed performance^22^. Longitudinal analyses conducted using the Health and Retirement Study in a diverse, nationally representative sample of older adults found that greater perceived discrimination predicted reductions in executive function, processing speed, and visuospatial construction performance^27^. Research conducted through the Black Women’s Health Study found that among older Black women, greater self-reported perceived discrimination predicted lower subjective memory for those who experienced more discrimination^28^.

A rapidly growing literature has turned its focus to investigate how perceived discrimination influences brain function and structure. In a cross-sectional lifespan study greater perceived discrimination predicted aberrant amygdala functional connectivity among a diverse sample^21^. Separately, during a task designed to assess neural responses to traumatic emotional images, more racial discrimination predicted greater functional activity in regions important for emotional regulation and attention among Black women^23^. Furthermore, independent of experiences of trauma and post-traumatic stress disorder, more perceived discrimination correlated with reduced fractional anisotropy, a measure of white matter structural integrity, in a number of white matter pathways among trauma-exposed Black women^24^. More recently, greater perceived discrimination was associated with lower hippocampal volume and greater white matter hyperintensities among older adults^25^. These findings expose the importance of characterizing the impact of perceived discrimination on neurocognitive integrity to determine how the sociocultural environment influences brain health. Altogether, the above findings provide ample evidence for the negative impact of perceived discrimination on neurocognitive health.

Complementing this literature, studies investigating moderators of these relationships are critical to understanding targets for intervention. Sense of control, a correlate of self-efficacy, moderated physical and psychological health^29–31^. Sense of control, measured using the Midlife Development Inventory (MIDI) Sense of Control scale, can be divided into two independent constructs, perceived constraints and personal mastery^30^. Personal mastery (i.e., internally perceived control over an event) and perceived constraints (i.e., externally perceived imposed obstacles) predicts well-being^30, 31^. Research conducted by Pruessner and colleagues (2005), demonstrated that among younger and older adults, greater sense of control predicted larger hippocampal volume^32^. Moreover, in older adults, sense of control attenuated the impact of physiological stress on both global brain volume and cognitive decline^32^. Here, we focus on personal mastery and adapt its definition by defining it as an individual’s self-perceived agency over their everyday life. It has been previously reported that greater, versus lower, personal mastery enables individuals to extricate themselves from unsolvable tasks^33^. In a sample of older African American and Afro-Caribbean adults, personal mastery partially explained how perceived discrimination correlated with psychological distress suggesting that understanding the impact of personal mastery is critical in the study of perceived discrimination on psychological distress^34^. Considered altogether, the perceived discrimination and sense of control literature provides evidence for a direct impact of psychosocial stress on brain health and suggests a moderating role of personal mastery on brain structure.

The current study’s goal was to investigate the relationship between perceived discrimination and MTL brain structures that are vulnerable to chronic stress and important for stress and emotional regulation. Thus, based on the extant literature we hypothesized that greater perceived discrimination would predict lower amygdala and anterior hippocampal volume. We also hypothesized that greater personal mastery would attenuate the aforementioned relationship. Here, we provide evidence for the relationship between psychosocial stress, perceived discrimination, and smaller brain volume, and a moderating role of personal mastery on these relationships. Our results complement a quickly growing literature on the relationship between perceived discrimination and neurocognitive health and provide evidence of the beneficial role of personal agency.

## Materials and Methods

### Participants

Data for this study was compiled from two pilot projects (study 1: Alzheimer’s Association Research Grant Chronic Stress and Aging Study; study 2: Boston University Alzheimer’s Disease Center (ADC) Pilot Grant) investigating the impact of experiences of perceived discrimination on brain structure in older adults (n = 36, 55 – 86 years; 58% women). Participants from Study 1 were recruited from the greater Boston area via flyers and advertisements in local papers. Participants from Study 2 were recruited through the Boston University ADC Health Outreach Program for the Elderly (HOPE) Study. Data collection began in 2018 and was ended in March 2020 due to the COVID-19 pandemic.

Study 1: Inclusion criteria included being between 50 to 80 years of age, identifying as non-Latinx Black/African (Black) American or White/European (White) American, fluent in English, a non-smoker, and a resident of the Greater Boston Area. Participants were excluded if they had any major signs or symptoms suggestive of neurological or psychiatric conditions, or disorders that are known to affect the medial temporal hippocampal system (e.g., epilepsy, clinical diagnosis of depression, post-traumatic stress disorder, etc.), or conditions that affect HPA axis function (e.g., Cushing’s disease).

Study 2: Inclusion criteria included being between 50 and 85 years of age, identify as non-Hispanic Black or White Americans, an ADC research diagnosis of “Control” (i.e., cognitively unimpaired), available MRI data (a T1-weighted structural scan), and fluent in English. Eligible participants were contacted by the ADC staff to determine interest in this research. Similar to Study 1, participants were excluded if they had any major signs or symptoms suggestive of neurological or psychiatric conditions, or disorders known to affect the medial temporal hippocampal system (e.g., epilepsy, clinical diagnosis of depression, post-traumatic stress disorder, etc.), or conditions that affect HPA axis function (e.g., Cushing’s disease).

All participants provided informed, written consent using procedures approved by the Boston Medical Campus Institutional Review Board, and this research was conducted under the guidelines of the Declaration of Helsinki. Data is available upon reasonable request and upon establishment of a formal data sharing agreement.

### Experiences of Discrimination

The Williams Major Discrimination questionnaire is a survey tool that has been extensively used in epidemiological studies to assess subjective interpersonal, perceived social discrimination^15^ across the lifetime. Respondents are posed a series of 9 questions related to everyday situations and asked to indicate whether they experienced discrimination and subsequently assign the reason for which they were discriminated against (e.g., sex, race, nationality, class, sexual orientation, etc.). Sample questions include: *At any time in your life, have you ever been unfairly fired; Have you ever been unfairly stopped, searched, questioned, physically threatened or abused by the police*? Following the work of Barnes and colleagues (2012), scores were summed for a minimum score of 0 and a maximum score of 9^22^. Higher scores reflect more experiences of discrimination.

### Perceived Stress Scale

We used the Perceived Stress Scale (PSS)^35^ to control for potential confounds of everyday life stress on stress due to perceived social discrimination. The PSS is a 10-item questionnaire that assesses how stressful an individual perceived their life to be over the course of the previous month. Participants were asked to select how much they agreed with each statement using a 5-point Likert scale ranging from *Never* to *Very Often*. Some example questions included: “In the last month, how often have you been upset because of something that happened unexpectedly?*”; “*In the last month, how often have you felt difficulties were piling up so high that you could not overcome them?*”*. PSS scores were calculated as the averaged sum of all items after reverse-scoring positively phrased items. Higher scores indicate greater perceived stress.

### Sense of Control

We used the MIDI Sense of Control questionnaire, a 12-item scale, to investigate the moderating influence of sense of control^29, 30^. Specifically, we used the personal mastery subscale to determine whether respondents self-perceived agency in responding to their experiences. Questions included items such as: *I can do just about anything I set my mind to, Other people determine most of what I can and cannot do*. Personal mastery was used as a proxy for personal agency. Higher scores reflect greater personal mastery.

## Magnetic Resonance Image Acquisition and Image Analysis

### MRI Acquisition

Participants from both projects were scanned at the Boston University Chobanian & Avedisian School of Medicine Center for Biomedical Imaging using the same Alzheimer’s Disease Neuroimaging Initiative pulse sequences, collected on a 3T Philips Achieva scanner with an 8-channel head coil. We obtained high-resolution T1-weighted structural scans (multi-planar rapidly acquired gradient echo images; SENSitivity Encoding P reduction: 1.5, S reduction: 2; TR = 6.7 ms, TE = 3.1 ms, flip angle = 9°, field of view = 25 cm, Matrix Size = 256 × 254, 150 slices, resolution = 0.98 mm × 0.98 mm × 1.22 mm).

### Regions of Interest

We conducted all automatic segmentations using FreeSurfer 7.1.0, a well-documented and free software available for download online (http://surfer.nmr.mgh.harvard.edu/)^36^. In brief, FreeSurfer is a standardized, automatic segmentation tool that probabilistically approximates subcortical and archicortical volume based on postmortem brain structure analyses^37^. Pre-processing includes motion correction and averaging^38^, non-brain tissue removal^39^, intensity normalization^40^, gray matter white matter boundary tessellation, Talairach transformation, and segmentation of white matter and deep gray matter volume^36, 41^. We used the *recon-all* command to obtain hippocampus and amygdala volumes. We also obtained hippocampal subfield volume which were generated using a probabilistic tool based on ultra-high-resolution *ex vivo* MRI data^42^. This also provided delineations of anterior and posterior hippocampus^42^. We had no hypothesis regarding laterality thus regions were assessed bilaterally. Finally, we used estimated intracranial volume (EICV), which was calculated based on the transformation of each individual participant’s T1 volume to the atlas template space for normalization^43^.

### Statistical Analyses

Statistical analyses were conducted using R (4.0.0) and RStudio (1.2.5042). Primary outcome variables were tested for normality using the Shapiro-Wilk test, and were normally distributed. Continuous variables were summarized by mean, range, and standard deviation. Sex was summarized by percentage. For descriptive purposes, demographic characteristics were grouped by sex (men and women) and racial group (Black and White) and were analyzed with the Wilcoxon rank-sum test due to unequal sample sizes.

We used multiple regression analyses, controlling for sex, education, current stress, and EICV. Continuous predictor variables were standardized by mean-centering and scaling by 2 standard deviations^44^. Personal mastery and perceived constraints (grouped into three levels as the mean ±1 standard deviation) served as the moderating variables in the moderation analyses, calculated using the *Interact* package in R.. Statistical significance was set at *p_FDR_* < .05, using the false discovery rate (FDR) for multiple comparison correction. In our exploratory analyses (described below) we did not correct for multiple comparisons, but provided confidence intervals, *F* statistics, and adjusted*-R*^2^ for comparison in addition to *p*-values^45^.

## Results

### Participant Characteristics

Participant characteristics for the overall sample are described in Table 1. For descriptive purposes, participant characteristics are also listed by race and by sex. Wilcoxon rank-sum tests showed that Black participants were younger, had fewer years of education, endorsed greater perceived discrimination and slightly higher perceived stress. Personal mastery in men and women differed at the trend level with women endorsing greater personal mastery.

**Table 1.** Participant Demographics. Significant results are denoted by dagger and asterisks († < .10, * < .05, ** < .01)

| **Variables** | **Overall**<br>N = 36<br>M (SD) | **Black**<br>N = 11<br>M (SD) | **White**<br>N = 25<br>M (SD) | **Male**<br>N = 15<br>M (SD) | **Female**<br>N = 21<br>M (SD) |
| --- | --- | --- | --- | --- | --- |
| Age (years) | 70.8 (6.8) | 66.23 (8.3) | 72.8 (5.0) * | 69.2 (8.5) | 72.0 (5.3) |
| Education | 16.4 (2.1) | 14.8 (1.7) | 17.1 (2.0) ** | 16.4 (2.1) | 16.4 (2.2) |
| Personal<br>Mastery | 22.1 (5.7) | 20.1 (6.6) | 23.0 (5.2) | 20.7 (5.2) | 23.0 (6.0) † |
| Perceived<br>Constraints | 46.6 (10.1) | 45.6 (8.2) | 47.0 (10.9) | 45.5 (7.5) | 47.3 (11.7) |
| Perceived<br>Stress | 11.4 (8.3) | 16.6 (10.5) | 9.1 (6.0) † | 11.3 (8.1) | 11.5 (8.6) |
| Perceived<br>discrimination | 1.8 (1.8) | 2.8 (2.0) | 1.3 (1.5) * | 2.1 (2.3) | 1.5 (1.3) |

### Association between perceived discrimination and amygdala volume and anterior hippocampal volume

Using multiple regression analyses, we tested whether perceived discrimination correlated with bilateral amygdala and anterior hippocampal volumes, controlling for current stress, sex, education, age, and EICV. Higher perceived discrimination correlated with lower bilateral amygdala (*ß* = -180.43, CI [-323.21, -37.66], *t*(30) = -2.59, *p_FDR_* = .03; model: *F*(6,29) = 3.22, *R*^2^_*adj*_ = .28) and anterior hippocampal (*ß* = -148.93, CI [-297.85, -0.0006], *t*(30) = -2.05, *p_FDR_* = .05; model: *F*(6,29) = 2.22, *R*^2^_*adj*_ = .17) volume (see Figure 1). Perceived discrimination did not correlate with posterior hippocampus volume (*ß* = -60.80, CI [-147.81, 22.20], *t*(30) = -1.51, *p* = .14; model: *F*(6,29) = 3.14, *R*^2^_*adj*_ = .27). We also tested whether perceived discrimination predicted volume in our regions of interest stratified by racial group and by sex and found no significant correlations (see Supplementary Table 1).

**Figure 1.**
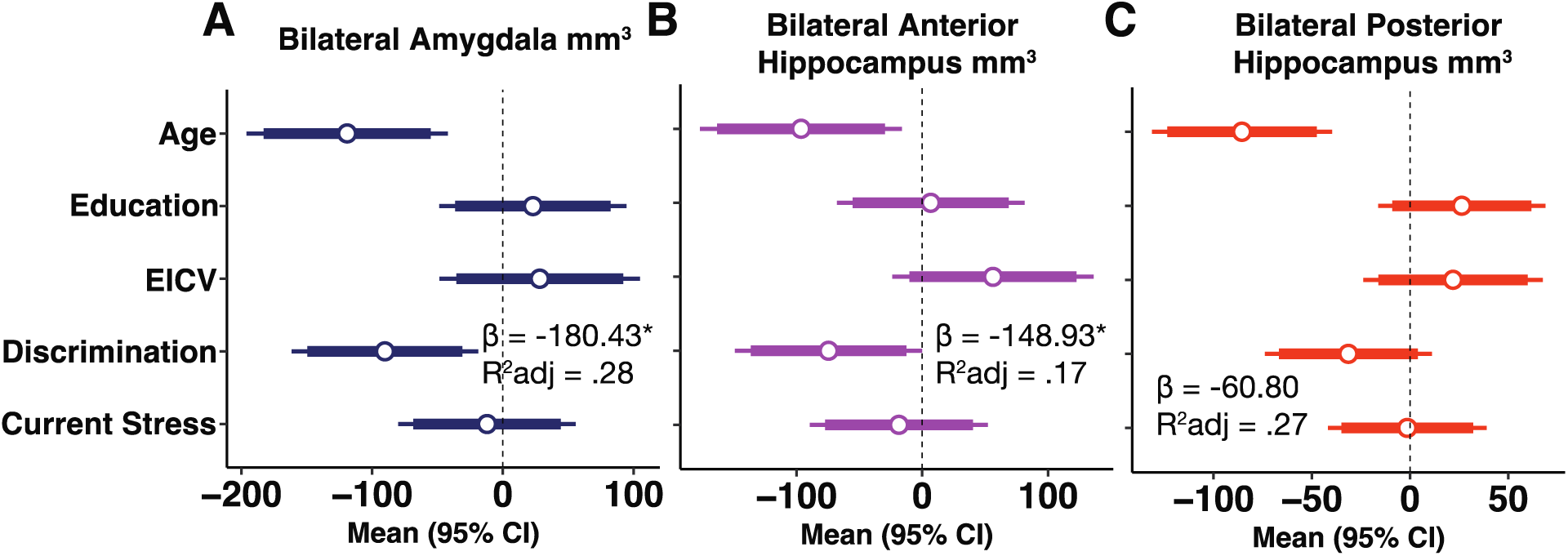
Forest plots representing multiple regression analyses with covariates, correlating perceived discrimination with A) bilateral amygdala volume, B) bilateral anterior hippocampus volume, and C) bilateral posterior hippocampus volume. Beta coefficients and adjusted-R^2^ are presented for each model, with 95% (inner 90%) confidence intervals. (*p < .05)

### Exploratory analyses by hemisphere and hippocampal subfield

We explored the relation of perceived discrimination to amygdala and anterior hippocampal volume by hemisphere. Perceived discrimination was associated with lower left but not right amygdala volume (see Table 2). Additionally, perceived discrimination did not correlate with left nor right anterior hippocampal volume (see Table 2).

**Table 2.** Exploratory analyses of amygdala and anterior hippocampal volume by hemisphere. Significant results are bolded (**p < .01)

| ROI | $\beta$ | 95% CI | $t(30)$ | $p$ | $F(6,29)$ | $R^2_{adj}$ |
| --- | --- | --- | --- | --- | --- | --- |
| Left Amygdala | **-199.46** | -344.69, 54.23 | -2.81 | <.01** | 3.56 | .31 |
| Right Amygdala | -161.41 | -337.11, 14.29 | -1.88 | .07 | 1.92 | .14 |
| Left Hippocampus | -170.77 | -355.00, 13.47 | -1.90 | .07 | 2.40 | .19 |
| Right Hippocampus | -127.08 | -285.68, 31.52 | -1.64 | .11 | 1.15 | .03 |

We further tested for the association between perceived discrimination and hippocampal subfield volume. Based on voxel size resolution, we combined the DG/CA3/CA4 subregions then tested for associations between perceived discrimination and bilateral anterior and posterior subfield volumes (DG/CA3/CA4, CA1, and subiculum). Greater perceived discrimination correlated with lower anterior but not posterior CA1 volume (see Table 3). There were no associations between perceived discrimination and anterior nor posterior DG/CA3/CA4 volume (see Table 3). Finally, we found that perceived discrimination negatively predicted anterior but not posterior subiculum (see Table 3).

**Table 3.** Exploratory analyses of bilateral hippocampal subregion volume. Significant results are bolded. (*p < .05, **p < .01)

| ROI | $\beta$ | 95% CI | $t(31)$ | $p$ | $F(5,30)$ | $R^2_{adj}$ |
| --- | --- | --- | --- | --- | --- | --- |
| Anterior CA1 | **-57.32** | -109.34, -5.30 | -2.25 | **.03\*** | 2.06 | .13 |
| Anterior DG/CA3/CA4 | -21.72 | -62.67, 19.23 | -1.08 | .29 | 1.79 | .10 |
| Anterior Subiculum | **-21.95** | -37.71, -6.19 | -2.84 | **<.01\*\*** | 2.29 | .16 |
| Posterior CA1 | -5.28 | -21.17, 10.60 | -.68 | .50 | 1.49 | .07 |
| Posterior DG/CA3/CA4 | -7.75 | -38.29, 22.79 | -.52 | .61 | 1.90 | .11 |
| Posterior Subiculum | -10.73 | -25.96, 4.49 | -1.44 | .16 | 4.50 | .33 |

Based on significant results in the anterior hippocampus, we next explored the relationship between perceived discrimination and anterior hippocampal subfields by hemisphere. Perceived discrimination significantly correlated with left anterior CA1 (*ß* = -65.79, CI [-130.12, -1.45], *t*(31) = -2.09, *p* = .05, model: *F*(5,30) = 1.95, *R*^2^_adj_ = .12) and subiculum volume (*ß* = -29.00, CI [-51.13, -6.88], *t*(31) = -2.68, *p* = .01, model: *F*(5,30) = 2.15, *R*^2^_adj_ = .14). Perceived discrimination did not significantly predict right anterior CA1 or subiculum, nor left nor right DG/CA3/CA4 volume (see Supplemental Table 2).

### Personal mastery attenuates the relationship between perceived discrimination and anterior hippocampal and amygdala volumes

We conducted moderation analyses separately by personal mastery and perceived constraints since they are purported to test independent effects of sense of control^31^. Personal mastery, but not perceived constraints (see Supplemental Table 3), interacted with perceived discrimination to predict bilateral amygdala (*ß* = 450.96, CI [5.43, 896.48], *t*(29) = 2.07, *p* = .05; model: *F*(7,28) = 3.55, , *R*^2^_adj_ = .34) and anterior hippocampus (*ß* = 498.20, CI [27.38, 969.02], *t*(29) = 2.17, *p* = .04; model: *F*(7,28) = 2.44, *R*^2^_adj_ = .22) volume, such that personal mastery attenuated the relationship between discrimination and regional volumes (see Figures 2, 3). We conducted a post-hoc test of simple slopes for each region of interest in Table 4.

**Figure 2.**
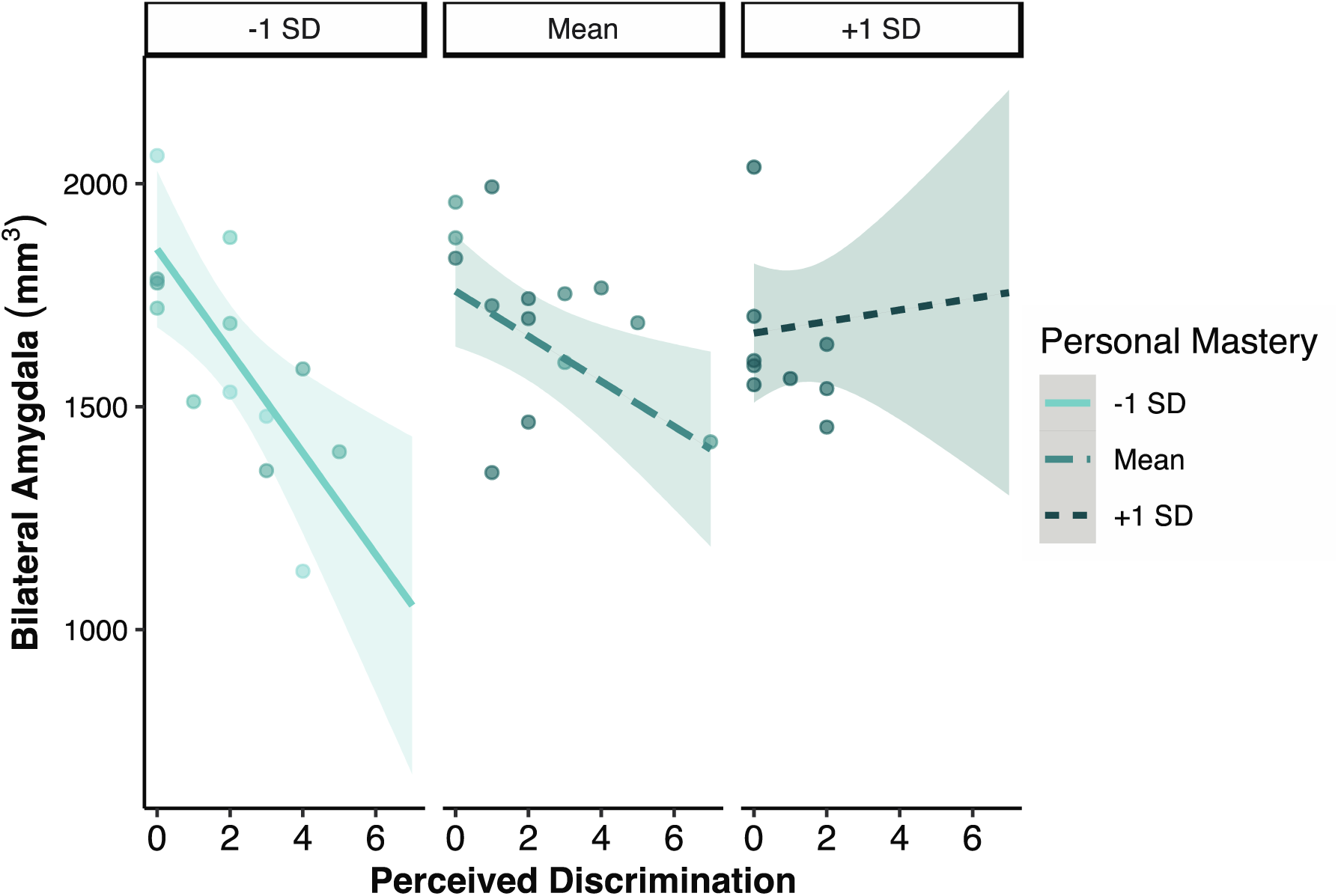
Perceived discrimination and personal mastery interacted to predict bilateral amygdala volume such that as personal mastery increased, the relationship between perceived discrimination and bilateral amygdala volume was attenuated.

**Figure 3.**
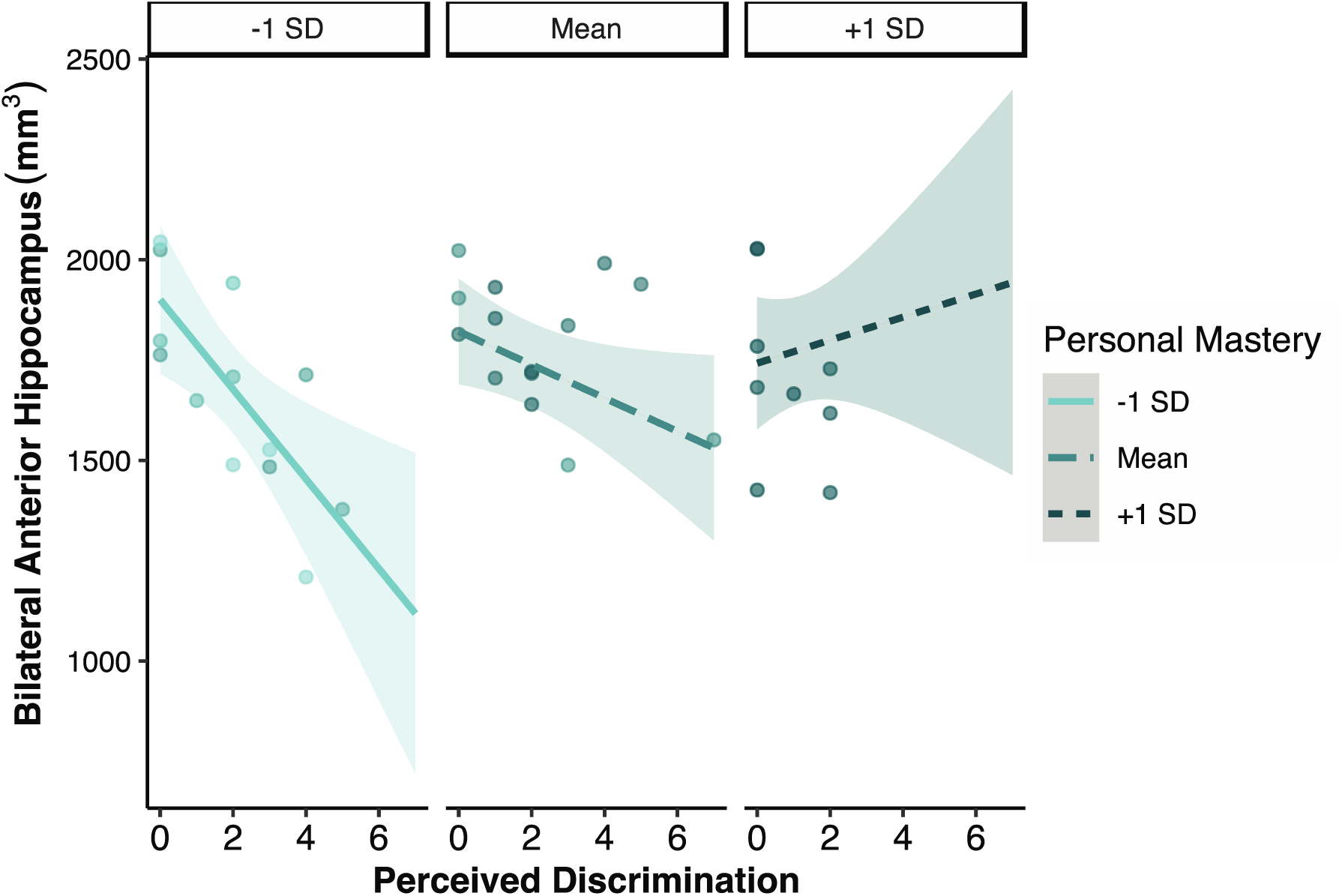
Perceived discrimination and personal mastery interacted to predict bilateral amygdala volume such that as personal mastery increased, the relationship between perceived discrimination and bilateral amygdala volume was attenuated.

**Table 4.** A test of simple slopes showed that the relationship between perceived discrimination and our regions of interest was attenuated at one standard deviation below (lower personal mastery – mean score of 16.35) and at the mean (average levels of personal mastery – mean score of 22.08) of personal mastery. \**p* ≤ .05, \*\**p* ≤ .01.

|  |  |  |  |
| --- | --- | --- | --- |
| Bilateral Amygdala | $\beta$ | $p$ | 95% CI |
| -1 *SD* | **-114.11** | < .01** | -186.16, -42.05 |
| At the mean | **-50.58** | .01** | -89.17, -11.99 |
| +1 *SD* | 12.95 | .73 | -62.32, 88.22 |
| Bilateral Anterior Hippocampus | $\beta$ | $p$ | 95% CI |
| -1 *SD* | **-111.65** | .01** | -187.80, -35.51 |
| At the mean | **-41.47** | .05* | -82.25, -.69 |
| +1 *SD* | 28.72 | .47 | -50.82, 108.26 |

## Discussion

This study’s primary objective was to investigate whether perceived discrimination, a salient psychosocial stressor, predicted amygdala and anterior hippocampal volume among older adults. We also sought to understand the moderating role of personal mastery on these relationships. We tested the hypothesis that higher perceived discrimination would correlate with lower amygdala and anterior hippocampal volume. Moreover, we tested whether higher levels of personal mastery would attenuate these relationships. We determined that more experiences of perceived discrimination correlated with lower amygdala and anterior hippocampal volume. We then demonstrated that higher personal mastery attenuated these relationships. We conducted exploratory analyses examining these regions by hemisphere and by hippocampal subfields. Greater perceived discrimination correlated only with left amygdala volume. Additional exploratory analyses showed greater perceived discrimination correlated with smaller anterior CA1and subiculum volume. Moreover, when we explored these associations by hemisphere, these associations were lateralized to the left hemisphere.

### Perceived discrimination is correlated with amygdala and anterior hippocampal volume

The amygdala and anterior hippocampus are well-characterized in their role in emotional and stress regulation^2,46^. We thus hypothesized that these regions may be particularly sensitive to the impact of perceived discrimination, a chronic psychosocial stressor. We specifically limited our focus to the amygdala and anterior hippocampus given their roles in modulating hypothalamic-pituitary-adrenal axis function, responsible for modulating the stress response, and their recruitment in emotional regulation^4,47^. Although longitudinal research is needed to measure causality or directionality between perceived discrimination and human neurocognition, we can use the extant literature to investigate how perceived discrimination impacts brain structure. It is well-established in animal models of stress that these brain regions undergo structural reorganization during stressful events^2,48^. For example, among tree shrews exposure to a dominant conspecific resulted in dendritic atrophy in the hippocampal CA3^1^ and psychosocial stress exposure impaired hippocampus-dependent memory and was associated with smaller hippocampal volume^49^. In comparison, among humans, greater overall stress predicted lower hippocampal volume^11^. More recently, perceived discrimination predicted aberrant amygdala functional connectivity^21^ and reductions in hippocampal volume^25^. Adding to this literature, we found that perceived discrimination negatively correlated with amygdala volume and had a localized relationship with the anterior hippocampus. In our supplemental analyses we found that when stratified by racial group and sex, perceived discrimination did not correlate with amygdala and anterior hippocampal volume. Despite differences in the stress response by sex we found no sex effect of perceived discrimination^2, 58^.

Inflammation provides one potential neurobiological pathway by which we may see a deleterious impact of perceived discrimination on neurocognitive health. This may include changes to underlying neural architecture and cognitive ability. Greater perceived discrimination has been associated with greater inflammation in humans^18, 50–54^. Greater inflammation, in turn, has also been shown to compromise cognition in older adults^52, 55^. Complementing this research, in rodents, inducing an inflammatory response in the hippocampus has been shown to impair hippocampal-dependent function^56, 57^. Together, this suggests a role for inflammation as a neurobiological mechanism underlying the link between perceived discrimination and neurocognitive health. Seminal work conducted by Geronimus and colleagues (2006) found that Black Americans showed greater allostatic load, a composite physiological measure of biological dysregulation in response to chronic stress, compared to White Americans^59^. When the groups were broken down by both race and sex, there was a significant difference whereby Black women showed the highest allostatic load across the lifespan, followed by Black men, White women, and White men. Allostatic load has been examined as a mechanism underlying the impact of chronic stress on health, including its influence on the plasticity of the human brain^3, 60^. Future research should collect physiological measures of allostatic load, including measures of inflammation, in order to determine whether allostatic load may mediate the relationship between perceived discrimination and brain structure in aging.

Finally, for our exploratory analyses, we anticipated that there would be a primary effect of perceived discrimination on the DG/CA3/CA4 due to its neuroplastic role in the production and maintenance of adult-born neurons and its sensitivity to stress^12, 61^. However, greater perceived discrimination correlated with lower CA1 and subiculum volume. The hippocampal CA1 and subiculum subregions are engaged in the encoding and retrieval of episodic memories^62, 63^. In healthy aging we see a reduction of both volume and neurons in the CA1 and subiculum in humans, and this effect is increased in Alzheimer’s disease^64^. Our exploratory results suggest that the CA1 and subiculum may be negatively impacted by perceived discrimination and provide a novel avenue of research on the sensitivity of hippocampal subfields in large-scale research on perceived discrimination. Additionally, research conducted within the context of perceived discrimination should use tasks that tax the function of these dissociable regions in order to confirm conjectures of functional impairment associated with greater perceived discrimination.

### Personal mastery attenuates the relationship between perceived discrimination and amygdala and anterior hippocampal volume

Although these neurobiological mechanisms play a part in explaining the harmful effects of perceived discrimination, it does not fully explain individual differences in how a person may respond to a stressful experience, or the agency they feel during this experience. A framework developed by Fani and Khalsa (2022), may explain, in part, some of these differences^65^. They hypothesize that in those experiencing racial discrimination, there may be a disruption of bodily and cognitive function as a result of experiencing and perceiving discrimination. Here, we found that as personal mastery increased, the relationships observed between perceived discrimination and amygdala and anterior hippocampus was negated. The amygdala and hippocampus are implicated in a number of disease states, including depression and may be disrupted as a result of rumination^66, 67^. We hypothesize that personal mastery may interrupt greater rumination via feeling greater agency during the discriminatory experience. Despite our results, the role of personal mastery may be more complicated depending on previous life experience^68^. In a study of childhood trauma and emotional reactivity to daily events, in participants who experienced childhood trauma, greater personal mastery predicted lower perceived sense of well-being^68^. Altogether, these results suggest a more intricate model of the impact of personal mastery on brain health, and this may be complicated by experiences earlier in life.

## Limitations

Despite our *a priori* focus on the MTL, a potential limitation of the study, and a subject for future investigation, is to study regions of interest outside of the MTL additionally recruited in stress and emotional regulation^2^. For example, the ventromedial prefrontal cortex, has been posited to inhibit amygdala function in stress and emotional dysregulation^69–71^. A second limitation is that our approach to understanding perceived discrimination is coarse in that we ask participants to retrospectively report experiences of discrimination and did not inquire about frequency of events, which may induce potential recall bias into our study. Future studies should consider strategies such as ecological momentary assessment which provide more comprehensive, real-time metrics to evaluate experiences of discrimination^73^, or group-specific studies that focus on experiences of discriminatory factors that do not depend on conscious awareness^74^. Due to our smaller sample size, we were unable to parse out differences related to different identities (e.g., race, socioeconomic status, gender) or their intersection^72^ which may individually or interactively contribute to our findings. We were also unable to determine whether personal mastery was driven by one group. This could have critical implications for translation: if data suggest that these effects occur only in one gender/race (i.e., one needs to be male or white in order to gain benefits from personal mastery), it might be counterproductive to tell marginalized people to take more control of their experiences if 1) they do not receive stress-related impacts neurocognition, and 2) what they really need is to experience less discrimination. Finally, while we characterize the relationship between perceived discrimination and brain volume, it is critical to assess cognitive and emotional function dependent on these brain regions to determine whether there is a corresponding negative impact of perceived discrimination on factors such as learning and memory performance, as has been seen for global cognition^22^.

Characterizing how perceived discrimination impacts brain health is imperative to health equity. In a rapidly aging population, it is integral that we understand the ways in which historical experiences of racism, misogyny, ableism, and other forms of discrimination impact brain health and well-being. Without an understanding of how the sociocultural environment “gets under the skin” to influence brain health we are unable to create interventions that may help ameliorate the impact of perceived discrimination, including interventions through health and public policy.

## Acknowledgements

This work was supported in part by the Alzheimer’s Association Research Grant (KS), a Robert Wood Johnson Foundation Health Policy Research Scholars grant (MAR), an NIH F99 Training Grant (MAR) (1F99NS124143-01A1), and the Boston University Clinical and Translational Science Institute (UL-TR000157). A portion of these data were previously reported and presented (https://doi.org/10.1002/alz.045394). We would also like to thank our participants for engaging in this research with us.

## Contributions

K.S., Y.C., and M.A.R. designed the study and developed the methodology. M.A.R. analyzed the data and wrote the first draft of the manuscript. M.A.R., R.A. A.E., and A.A. collected the data. K.S., Y.C., A.B., U.C., R.A., A.E., and A.A. provided critical feedback for revision of the manuscript.

## Competing interests

The author(s) declare no competing interests or conflicts of interest.

## Data Availability Statement

Data available upon reasonable request and upon establishment of a formal data sharing agreement.

## Notes

### Competing Interest Statement

The authors have declared no competing interest.

