## Supplemental Tables 1, 2, and 3 for "Personal Mastery Attenuates the Association between Greater Perceived Discrimination and Lower Amygdala and Anterior Hippocampal Volume in a Diverse Sample of Older Adults"

| | Amygdala | Anterior Hippocampus | Posterior Hippocampus |
|---|---|---|---|
| PD*Sex | 13.24 | -41.36 | -25.14 |
| | [-130.48, 156.97] | [-190.52, 107.79] | [-110.21, 59.93] |
| N | 36 | 36 | 36 |
| $R^2$ | 0.40 | 0.32 | 0.40 |
| PD*Group | -58.96 | -51.03 | -4.18 |
| | [-196.13, 78.20] | [-183.01, 80.95] | [-85.21, 76.84] |
| N | 36 | 36 | 36 |
| $R^2$ | 0.56 | 0.57 | 0.56 |

**Supplemental Table 1.** Interaction analysis of PD and sex. (Beta estimate [CI]; PD: perceived discrimination)

| | Left CA1 | Right CA1 | Left DG/CA3/CA4 | Right DG/CA3/CA4 | Right Subiculum |
|---|---|---|---|---|---|
| PD | **-32.89 *** | -24.43 | -9.04 | -12.68 | -7.45 |
| | CI [-65.06, -0.73] | CI [-53.42, 4.56] | CI [-32.59, 14.50] | CI [-34.75, 9.40] | CI [-15.45, .56] |
| N | 36 | 36 | 36 | 36 | 36 |
| $R^2$ | 0.25 | 0.17 | 0.20 | 0.20 | .14 |

**Supplemental Table 2.** Perceived discrimination and anterior hippocampal subfield by hemisphere, uncorrected. (Beta estimate [CI]; PD: perceived discrimination; $^*p \leq .05$)

| | Amygdala | Anterior Hippocampus |
|---|---|---|
| PD*Perceived Constraints | -12.52 | -49.03 |
| | CI [-88.34, 63.31] | CI [-128.00, 29.94] |
| N | 36 | 36 |
| $R^2$ | 0.39 | 0.31 |

**Supplemental Table 3.** Perceived constraint does not interact with PD to predict amygdala nor anterior hippocampal volume, uncorrected. (Beta estimate [CI]; PD: perceived discrimination)

Although we conducted an exploratory analysis of perceived constraints, we found no moderating effect of perceived constraints (see Supplemental Table 3). In contrast to personal mastery, we expected to see an inverse relationship such that at lower levels of perceived constraints, perceived discrimination would have a negligible influence on amygdala and anterior hippocampus volume. However, it is possible that perceived constraints and perceived discrimination may interact in other ways to influence neurocognitive integrity.
